# Pulmonary Hypertension Engine for Linked Experiments (PHELEX): a platform for the re-analysis of public transcriptomic data related to pulmonary hypertension in both animal models, and humans

**DOI:** 10.64898/2026.04.28.721394

**Authors:** Tanvi Nandani, Benjamin P Ott, Prusha Balaratnam, Stephen L Archer, Joshua Durbin, Charles C T Hindmarch

## Abstract

Pulmonary hypertension (PH) is a vasculopathy that results in elevated mean pulmonary arterial pressures over 20mmHg. Despite significant advances in research, PH still has a high mortality rate, and there is currently no cure for the disease. As with all biomedical fields, PH researchers have embraced the power of next generation technologies such as microarrays and RNA sequencing. Most of these data can be found on public repositories, which is usually a requirement for publication. While these repositories are rich sources of data, they require intermediate to advanced bioinformatics skills to access, download, and make these data useful. Here we present <u>**P**</u>ulmonary <u>**H**</u>ypertension <u>**E**</u>ngine for <u>**L**</u>inked <u>**E**</u>xperiments (<u>**PHELEX**</u>), which represents a comprehensive catalogue of all RNA sequencing data related to PH that is currently available on the Gene Expression Omnibus (GEO), hosted by the US National Centre for Biotechnology Information (NCBI). We identified 2,278 bulk RNA sequencing samples from human, mouse and rat, and built a searchable tool based on the metadata that is associated with each sample. PHELEX is a functional tool that allows selected studies to be highlighted, and parsed through Confidence, an analysis tool we have created, which will model the data based on user-defined classifiers, perform differential gene expression and pathway analysis, and present these data using standard graphics, and text-file results. PHELEX also allows PH researchers to cross-cut between discrete studies, facilitating de novo understanding of these data. As a robust searchable repository of genomic data, we hope that PHELEX will accelerate PH innovation and discovery, by allowing researchers to mine existing genomic data and thus better understand the molecular signatures that underpin PH.

## Introduction

Pulmonary hypertension (PH) is a lethal vasculopathy that impacts the lives of millions of people worldwide (*1*). Defined as a mean pulmonary artery (PA) pressure > 20mmHg, PH results in a poor prognosis for patients (*2*), and those suffering PH have a high mortality rate. There are five recognized PH groups: Group1 is defined as pre-pulmonary capillary pulmonary arterial hypertension (PAH) which may be idiopathic, heritable, or associated with connective tissue disease, portal hypertension, congenital heart disease, medications, or infections. Group2 is the most common form of PH and is related to left heart disease resulting in post-capillary (pulmonary venous) or combined pre and post capillary pulmonary hypertension. Group3 disease is secondary to underlying pulmonary disorders or environmental hypoxia and presents with a pre-capillary phenotype, Group4 disease is related to pulmonary artery obstructive disorders or chronic thromboembolic disease and presents with a pre-capillary phenotype, Group5 is a collection of mixed etiologies, including hemolytic anemias, myeloproliferative diseases and sarcoidosis, with both pre and post capillary phenotypes. Various animal models have been created to study different groups of PH, including (but not restricted to) the monocrotaline (MCT), and the Sugen Hypoxia (SuHx) rat models of Group1 PH, and other (*3*). Because most PH deaths can be attributed to right heart failure that results from sustained pulmonary pressures (*4*), much of the genomic research into mechanisms of Group 1 PH has been focused on the heart and lungs, as well as circulating cells from the blood. Indeed, most of the research has been focused on human patients (*5*), or on animal models (*3, 6-9*) created mostly in mice and rats. As with all biomedical fields, RNA sequencing has become a mainstay discovery tool with many groups profiling transcriptome responses in both preclinical animal models and in clinical cohorts to try and understand signature of genes that are regulated in multiple tissues.

The first twenty years of the genomic era have witnessed an unprecedented level of data collection estimated globally by the National Institutes of Health to be between 2-40 billion gigabytes of biological data. Importantly this rate of data collection is predicted to increase to 2-40 exabytes in the next decade. Much of this data is transcriptomic; describing whole genome expression levels in each study sample so that statistically robust differences in regulation of genes between conditions can be identified. Studies that employ omics (a suffix denoting comprehensive, system-level profiling such as transcriptomics (RNA), genomics (DNA), proteomics (protein), lipidomics (lipids), methylomics (methylome) etc., together with emerging omic domains yet to be fully understood) usually present a list of differentially regulated nodes and often attempt to place subsets of data into some biological context using tools such as Gene Ontology (GO) analysis or Gene Set Enrichment Analysis (GSEA). Most publications focus on a limited subset of genes that they prioritize for validation, leaving thousands of transcripts and GO pathways unexplored. One major problem that the omic revolution promised to address was the concept of the *ignorome*. A meta-analysis by Stoeger and colleagues has shown that the focus of most current biomedical research remains on the same 2000 genes that have been studied for the last three decades (*10*). This form of knowledge bias results in a small subset of genes that receive disproportionate funding and publication attention. While the cost of sequencing biological tissue at sufficient depth has decreased, we estimate the ‘per-sample’ cost of existing GEO data is $500-$1000 USD. As of 2024, GEO holds 6,500,000 individual samples (GSM), costing between $3.25 and $6.5 billion of publicly funded research dollars globally. This reflects over 200,000 studies, from over 6,000 organisms, with over 70,000 unique submitters, and corresponding to over 47,000 articles in PubMed Central (*11*). While these data have clearly been impactful, we suggest that the true benefits of these data reside beyond individual study analysis, and instead in the meta-analysis and crosscutting between studies. There is arguably more unmined genomic data than has been published, and so repurposing these existing transcriptomic data and metadata has the potential to greatly expand knowledge at minimal cost, provided the logistics of identifying and analyzing publicly accessible data sets can be overcome (*12*). We have anecdotally observed that majority of transcriptome papers, in multiple fields, rely on these comprehensive analyses to amplify the role of a small set of genes that are templates for validation, and sometimes functional analysis. This highlights the need for pathology specific databases that empower researchers to crosscut between studies and highlight novel targets.

Once transcriptomic or genomic data have been published in a paper, the authors are typically required (by their funder, the journal, or both) to upload the data to a public repository. The NIH funded National Centre for Biotechnology Information (NCBI) hosts transcriptomic data from every source, every species, and every paradigm within a repository called the Gene Expression Omnibus (GEO). GEO is part of the International Nucleotide Sequence Database Collaboration (INSDC), meaning that data submitted is continuously synchronized between three pillars; the NCBI, the European Bioinformatics Institute (EBI), and the DDBJ Sequence Read Archive (DRA) in Japan. This rich resource includes some built in functionality for analysis for certain datatypes so that standardized reanalysis of the data from individual papers can be performed. Experienced users can then identify, and extract these data based on keyword searches, together with their metadata, and analyze this on their own computers, or rely on the built-in GEO2R software (*11*) which can be used to analyze microarray data and some RNA sequencing datasets.

However, while utilization of these archived data is possible, we have identified significant barriers to discoveries that rely on data that has already been collected. These barriers include 1) requirement of intermediate to complex bioinformatics skills, 2) computational resources for storage and analysis of raw sequencing data, and 3) insufficient sample metadata (*14*). To overcome these challenges and help the PH community to accelerate research using this rich resource, here, we present our novel platform called **PHELEX**: **P**ulmonary **H**ypertension **E**ngine for **L**inked **Ex**periments (https://phelex.apps.meds.queensu.ca/). Focused on facilitating transcriptomic discovery relevant to pulmonary hypertension (PH), PHELEX has currently captured 106 datasets relevant to PH from GEO. Users can filter these datasets by species, tissue, sequencing modality, and other metadata, and then parse these outputs into *Confidence* (*13*), a turn-key application we have also created that analyzes counts files, performs differential gene expression analysis, and places prioritized genes into biological context with functional analysis. *Confidence* is designed to look for intersections between the outcome of the 4 key publicly available analysis tools (DESeq2, Limma, NOISeq and edgeR), providing a more robust understanding of the false discovery rate. While several platforms already exist that facilitate the analysis of these types of data, most of them only leverage a single analysis strategy, thus incurring the false discovery penalty and limiting the robustness of that platform. *Confidence* handles data differently because it parses counts data through 4 separate analyses (DESeq2(*14*), Limma(*15*), NOISeq(*16*) and edgeR(*17*)), providing a weighted score that maximises the user’s confidence in prioritized target genes.

With *Confidence* integrated into PHELEX, users can quickly identify RNA sequencing data related to PH, and associated meta-data, and then analyze these transcriptome data (aligned to the most recent species-specific genomes) using multiple pipelines so that they can reproduce and use existing public data. Moreover, PHELEX also allows for cross-cutting, (the process of comparing datapoints or variables between studies) resulting in a list of intersecting genes between discrete studies with similar experimental conditions and providing users with forest plots to show whether genes are moving in the same direction between studies. PHELEX has the potential to empower PH researchers without bioinformatics experience to mine, and re-mine public data and capitalize on the original investment meant to progress the research in our field.

## Methods

### 1) Application Overview

PHELEX is an integrated R Shiny application for exploration and analysis of PH-associated RNA-seq datasets from Gene Expression Omnibus (**Figure1**). It is composed of two interconnected components: (1) a local data retrieval and bioinformatics processing pipeline that compiles PH-relevant datasets from GEO, downloads raw sequencing data, and performs preprocessing, and (2) an online, interactive, frontend dashboard where researchers can explore, curate and analyze these datasets. Researchers can also perform an exploratory meta-analysis by comparing differential gene expression analysis results across datasets thereby identifying common differentially expressed genes.

**Figure 1:**
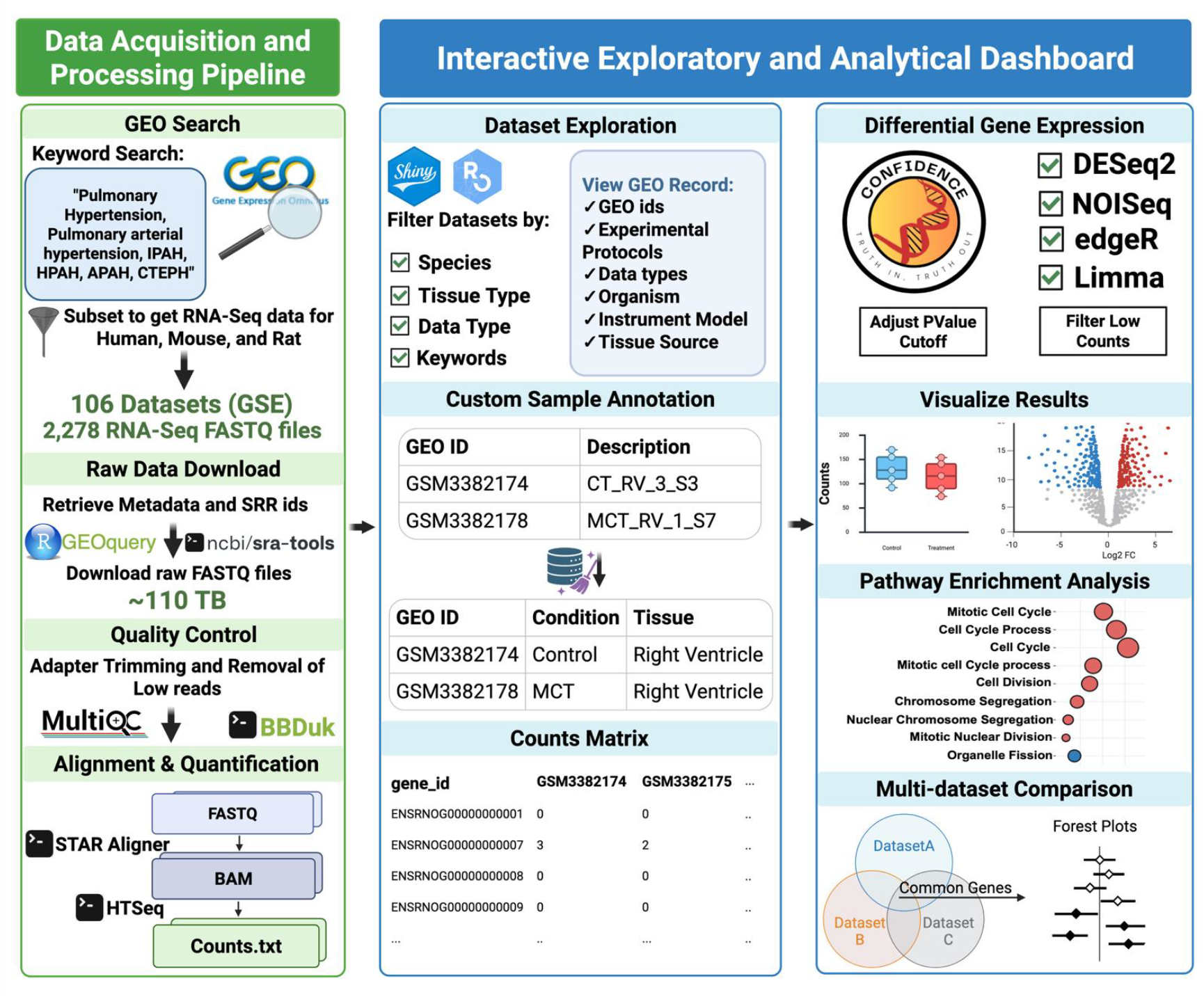
Overview of PHELEX Application: (Left) The data acquisition and processing pipeline searches GEO for PH-associated datasets, downloads raw RNA sequencing FASTQ files, and produces uniform count files through standardized quality control, alignment, and quantification. (Right) The interactive analytical dashboard enables researchers to explore datasets, curate sample metadata, perform differential gene expression analysis using four methods (DESeq2, edgeR, Limma, NOISeq) with *Confidence*, and compare results across multiple datasets.

As of the current release, the platform consists of counts files from 106 GSE Datasets consisting of 2,278 uniformly processed RNA sequencing files from *Homo Sapiens, Mus Musculus* and *Rattus Norvegicus* species, representing (to our knowledge) the largest systematically curated transcriptomic repository dedicated to PH research.

### 2) Data Retrieval and Curation

Records associated with all clinical groups of pulmonary hypertension were identified from the NCBI Gene Expression Omnibus on 13 March 2026 using the following keyword query:

(*((((“hypertension, pulmonary”[MeSH Terms] OR pulmonary hypertension[All Fields]) OR pulmonary arterial hypertension[All Fields]) OR idiopathic pulmonary arterial hypertension[All Fields]) OR (associated[All Fields] AND pulmonary arterial hypertension[All Fields])) OR (chronic[All Fields] AND thromboembolic[All Fields] AND (“hypertension, pulmonary”[MeSH Terms] OR pulmonary hypertension[All Fields]))) OR (hereditary[All Fields] AND pulmonary arterial hypertension[All Fields]*)

Metadata for all resulting records was compiled using GEOQuery (v3.22). In a secondary filtration step, the same PH-associated search terms were matched against study-level and sample-level metadata fields (title, summary, overall design, and sample characteristics). From this collection, 2,278 RNA sequencing samples from human, mouse and rat samples with available FASTQ files were identified.

Raw data files were downloaded from sequence read archive (SRA) using sra-tools (v3.4.1). The total size of retrieved transcriptomic sequencing data was approximately 110 TB. Raw data for other sequencing (scRNA, miRNA, ncRNA) and non-sequencing assay types (microarray, protein, and methylation profiling), and other species have been archived and will be incorporated in future releases of the platform to enable integrated multi-omics analysis.

### 3) Bulk RNA-seq data processing pipeline

Raw FASTQ files retrieved from SRA portal were processed through a standardized processing pipeline on Queen’s University’s high-performance computing cluster - Frontenac, running Rocky Linux (v8.10). Initial quality control was assessed for each sample using FastQC (v0.12.0). Adapter sequences, poly-A tails and low-quality bases were removed using BBDuk from the BBTools suite (v39.06). Adapter sequences were identified by kmer matching (matching short fixed-length nucleotide subsequences) and trimmed from the right end of reads (ktrim=r). For paired-end reads, a kmer length of 21 was used, with overlap-based adapter detection (tbo=t) and even trimming of both reads in a pair (tpe=t) enabled. For single end reads, a kmer length of 13 was used. For both data types, quality trimming was performed on both ends of reads (qtrim=rl) using a minimum quality score threshold of 15. Reads shorter than 30 bp after trimming were discarded. Post-trimming quality metrics were checked using MultiQC (v1.34).

Trimmed reads were aligned to respective reference genomes (*GRCh38*.*p14, GRCm39 and mRatBN7*.*2*) using the STAR (v2.7.11b) aligner tool. Reads were mapped allowing up to 20 multi-mapped locations, with a maximum mismatch-to-read-length ratio of 0.6. Alignments were filtered based on splice junction output and produced as coordinate-sorted BAM files, which were then indexed using SAMtools (v1.23.1). Gene-level read counts were quantified from the BAM files using HTSeq-count (v2.0.3) and using corresponding Ensembl gene annotations.

### 4) Dataset Exploration and Metadata Cleaning

Processed datasets within the PHELEX repository can be explored on the interactive RShiny dashboard (https://phelex.apps.meds.queensu.ca/). Datasets can be filtered by species, tissue source, and experiment type through the integrated Rentrez package (v1.2.4), which interfaces with the NCBI Entrez API to ensure raw sample metadata is displayed.

One of the challenges in reusing GEO data is the heterogeneity in data documentation in the sample metadata due to usage of open-ended, non-standardized textual description by the scientific community (*18*). To address this, PHELEX provides a metadata curation step in which researchers can select samples of interest and assign standardized condition labels (e.g., control, disease, treatment). Once samples have been selected and annotated, a gene-by-sample count matrix is assembled from the corresponding pre-processed counts files for further downstream differential gene expression analysis.

### 5) Differential Gene Expression, Gene Scoring and Enrichment Analysis with Confidence

Differentially expressed genes (DEG) from the selected datasets can be analyzed within the application through the integrated *Confidence* software (*13*) which utilizes DESeq2 (v1.48.2), Limma (v3.63.6), NOISeq (v2.52.0) and edgeR (v4.6.3). It also performs Principal component analysis (PCA) using the DESeq2 plotPCA. Volcano plots are generated on DEGs from each analytical package. Additionally, box plots can be generated for each DEG to visualize normalized gene counts between experimental conditions.

*Confidence* produces a combined analysis results table featuring results from each analysis package used. DEG in the results table also features a *Confidence* Score, which is a tally of the number of packages where a significant DEG is demonstrated to be up-or down-regulated (*13*). Based on a Confidence score threshold, the list of DEGs can further enriched using GOSt function from the gprofiler (v0.2.3) to perform pathway enrichment analysis using multiple databases simultaneously.

### 6) Multi-Dataset Comparison

Exploratory meta-analysis of two or more datasets can be performed within PHELEX. DEGs from each dataset can be filtered using a minimum *Confidence* score threshold, and overlapping genes across datasets can be analyzed with upset plots (UpSetR v1.4.0). For each overlapping gene, the direction of the regulation is determined from the log fold change values obtained from the analytical packages of *Confidence*. Genes of interest can be visualized using a forest plot displaying log2 fold-change estimates and 95% confidence intervals derived from DESeq2 (log2FC ± 1.96 × lfcSE).

## Results

### Summary of PH associated multi-omics data on Gene Expression Omnibus

The search on GEO using PH-associated terms returned 1,273 records comprising 334 submitter-supplied datasets (GSE), 21 curated datasets (GDS), 904 sample records (GSM), and 13 platform records (GPL). After filtering for PH-specific data, the results were to 319 series (GSE), 16 curated datasets (GDS), and 6,263 individual samples (GSM).

Systematic analysis of the curated PH-associated records revealed a diverse collection of data types, with RNA sequencing (n = 2,854) and microarray (n = 2,296) samples representing the most prevalent experiment types, followed by protein profiling, miRNA sequencing, and other sequencing modalities (**Figure 2a**). Most samples were derived from Homo sapiens (n = 4,342), followed by Mus musculus (n = 962) and Rattus norvegicus (n = 819), with additional samples from Ovis aries, Bos taurus, Peromyscus maniculatus, and Gallus gallus (**Figure 2b**). A clear shift from microarray to next-generation sequencing (NGS) is evident from 2016 onward, with NGS now representing the dominant data type across PH-associated datasets (**Figure 2c**).

**Figure 2:**
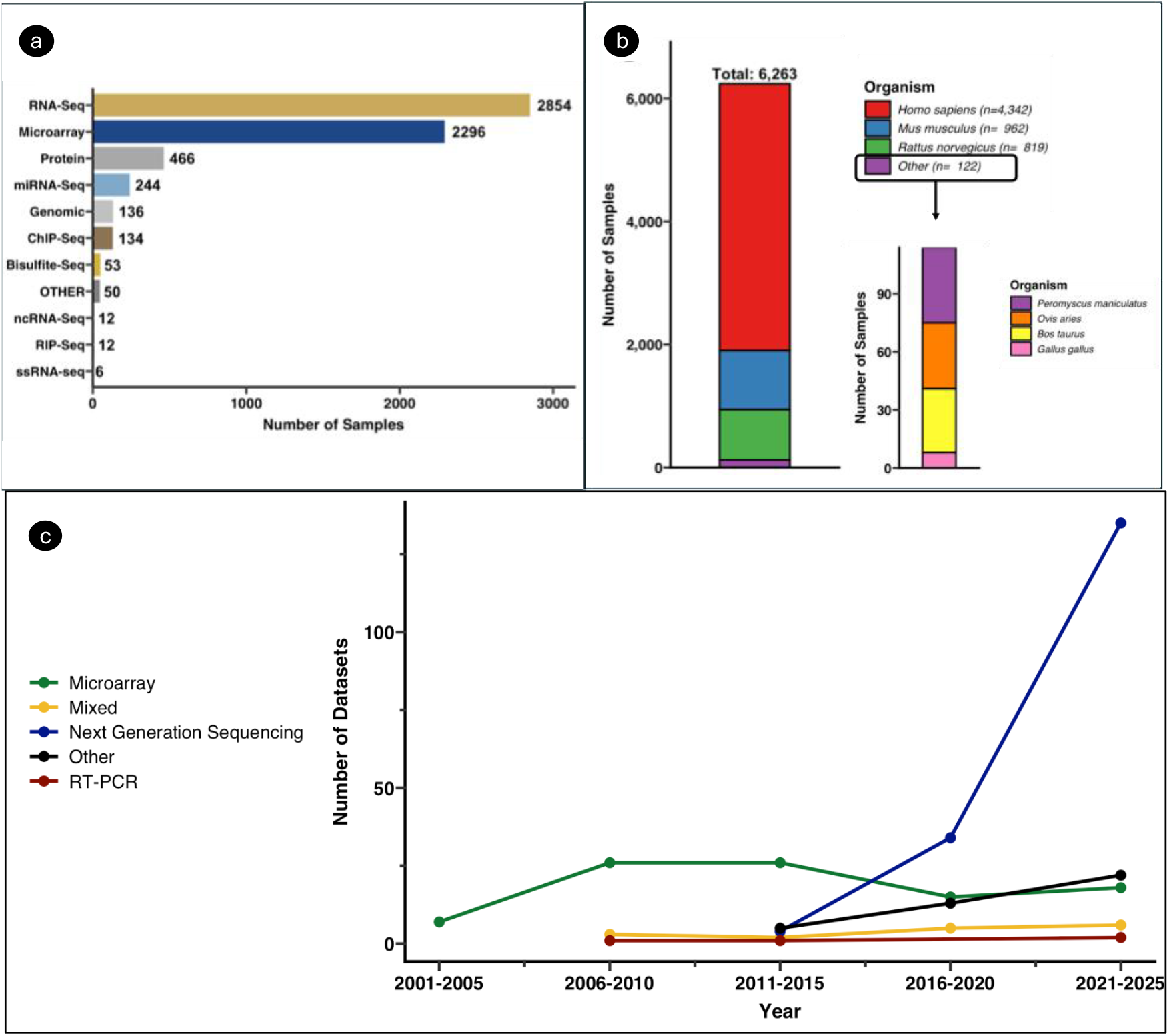
Summary of PH-associated data on GEO: (a) Distribution of samples by assay type, with RNA-seq and microarray as the most common modalities. (b) Distribution of 6,263 samples by organism as of March 2026. The majority are from Homo sapiens (n = 4,342), followed by Mus musculus (n = 962) and Rattus norvegicus (n = 819), with additional samples from Peromyscus maniculatus, Ovis aries, Bos taurus, and Gallus gallus. (c) Number of PH-associated dataset submissions to GEO by platform type over time, showing a sharp increase in next-generation sequencing datasets over the past decade.

Reanalysis of publicly available datasets from GEO requires extensive metadata curation, computational resources, and significant bioinformatics expertise, making it a time-consuming and often inaccessible process for many researchers. PHELEX was designed to address these challenges by providing a web-based platform that streamlines the reanalysis workflow, from dataset exploration and metadata curation to differential gene expression and pathway enrichment analysis.

PHELEX’s interactive analytical dashboard currently provides processed count matrices from 106 GSE datasets, comprising 1,425 human, 399 rat, and 455 mouse samples. The dashboard is implemented in R using the Shiny framework and is publicly accessible at: https://phelex.apps.meds.queensu.ca/.

To demonstrate the key functionalities of the platform, we analyzed samples from GSE240923 (*19*), a publicly available dataset (**Figure 3; Supplementary data**). This dataset is representative of the complexity typically encountered in PH-associated GEO datasets.

**Figure 3:**
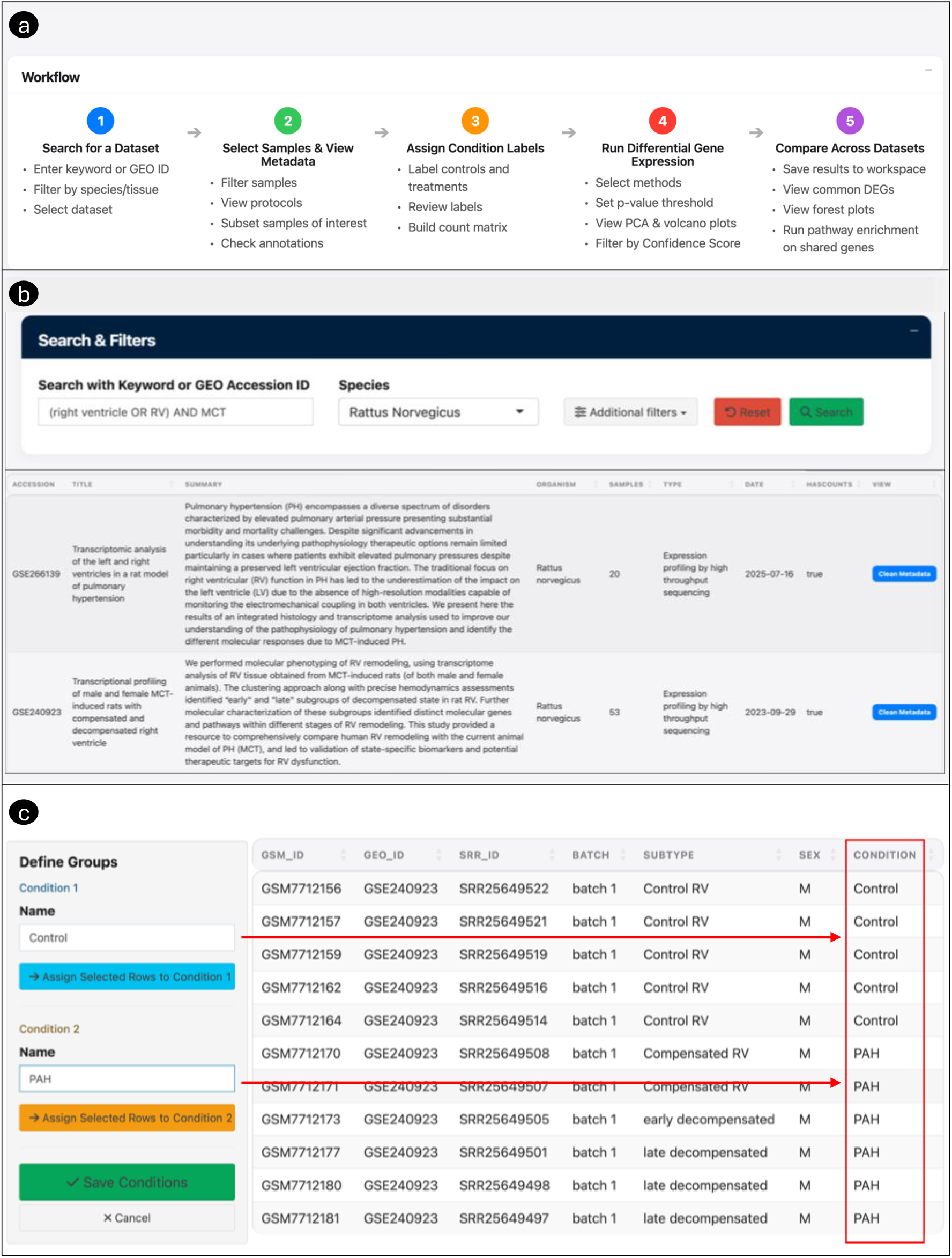
Exploring and curating datasets in PHELEX: a) The PHELEX workflow (b) The search module supports free-text keyword and GEO accession queries with filtering by species and dataset type. Search results display study metadata including title, summary, organism, sample count, and data type. (c) The metadata curation module allows users to assign custom condition labels (e.g., Control, PAH) to define experimental groups for downstream analysis

### 1) Dataset search and selection

Preprocessed datasets can be searched with keywords and filtered by species and dataset type (**Figure 3b**). Free-text Boolean search and direct GEO ID queries are also supported. In addition to preprocessed datasets, metadata for unprocessed PH-associated records can be browsed to identify datasets of interest. For example, a search for right ventricular (RV) tissue samples from monocrotaline (MCT) models returned four preprocessed datasets matching those criteria on PHELEX.

### 2) Clean Metadata and Build Counts Matrix

Sample-level metadata for a selected dataset can be explored and filtered within the metadata curation module (**Figure 3c**). Researchers can assign custom condition labels (e.g., Control, Compensated, Treatment) to define experimental groups for each selected sample. For GSE240923 (*19*), samples were subsetted to include only those from a single experimental batch and re-annotated as either “Control” or “PAH” to define the comparison groups for downstream analysis (**Figure 3c**). Once sample labels have been assigned, a gene-by-sample count matrix is assembled with the corresponding annotations, ready for differential gene expression analysis within the Confidence module. Both the curated metadata and the assembled count matrix are available for download for external use.

### 3) Differential Gene Expression Analysis with Confidence software

The assembled count matrix and curated metadata are imported into the integrated *Confidence* module for differential gene expression analysis (**Figure 4**). *Confidence* provides optional low-count gene filtering and generates principal component analysis (PCA) plots coloured by selected experimental variables, enabling exploratory assessment of sample clustering across multiple annotations. For GSE240923 (*19*), PCA for condition subtypes showed separation among control RV, compensated RV, early decompensated, and late decompensated samples (**Figure 4a**), while PCA for the two main experimental conditions showed clear separation between control and PAH groups (**Figure 4b**).

**Figure 4:**
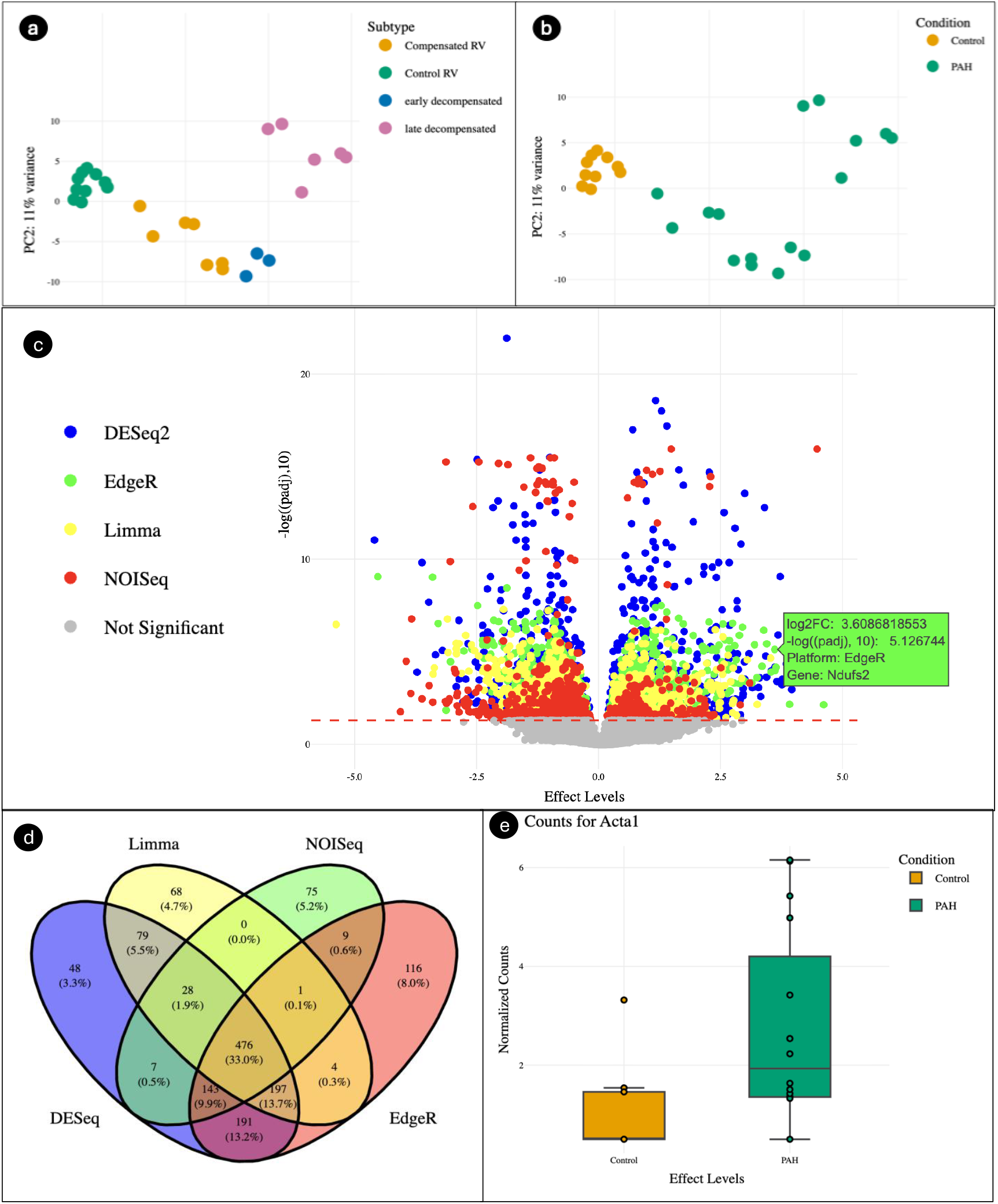
Differential gene expression analysis of GSE240923 with PHELEX. (a, b) Principal component analysis showing sample clustering by subtype and experimental condition. (c) Volcano plot displaying log2 fold-change and adjusted p-values for all tested genes, coloured by analytical method. (d) Four-way Venn diagram showing 476 genes identified as significantly differentially expressed by all four methods. (e) Box plot of normalized counts for Acta1.

Based on selected adjusted p-value threshold and the four available analytical methods (DESeq2, edgeR, limma, NOISeq), *Confidence* generates volcano plots displaying log2 fold-change and adjusted p-values for all tested genes, gene-level expression box plots for individual candidates, and a results table containing fold-change estimates, p-values, and *Confidence* Scores (**Figure 4c,e**). For GSE240923, differential gene expression analysis with all four methods identified 476 genes as significantly differentially expressed by all methods, as visualized in the four-way Venn diagram (**Figure 4d; Supplementary data**). Pathway enrichment analysis can subsequently be performed on the resulting differentially expressed gene lists, filtered by user-defined Confidence Score thresholds.

### 4) Multi-dataset Comparison

PHELEX enables cross-dataset comparison by allowing researchers to save differential gene expression results from multiple datasets to a session workspace. Saved results are compared using UpSet plots, which display the intersection of differentially expressed genes across datasets at user-defined Confidence Score thresholds (**Figure 5**). Forest plots are generated for shared genes, displaying DESeq2 log2 fold-change estimates with 95% confidence intervals (log2FC ± 1.96 × lfcSE) to assess the consistency of effect sizes and directionality across studies. Pathway enrichment analysis can then be performed on the resulting common gene set (**Figure 5c-f**). No user data are stored on the PHELEX server.

**Figure 5:**
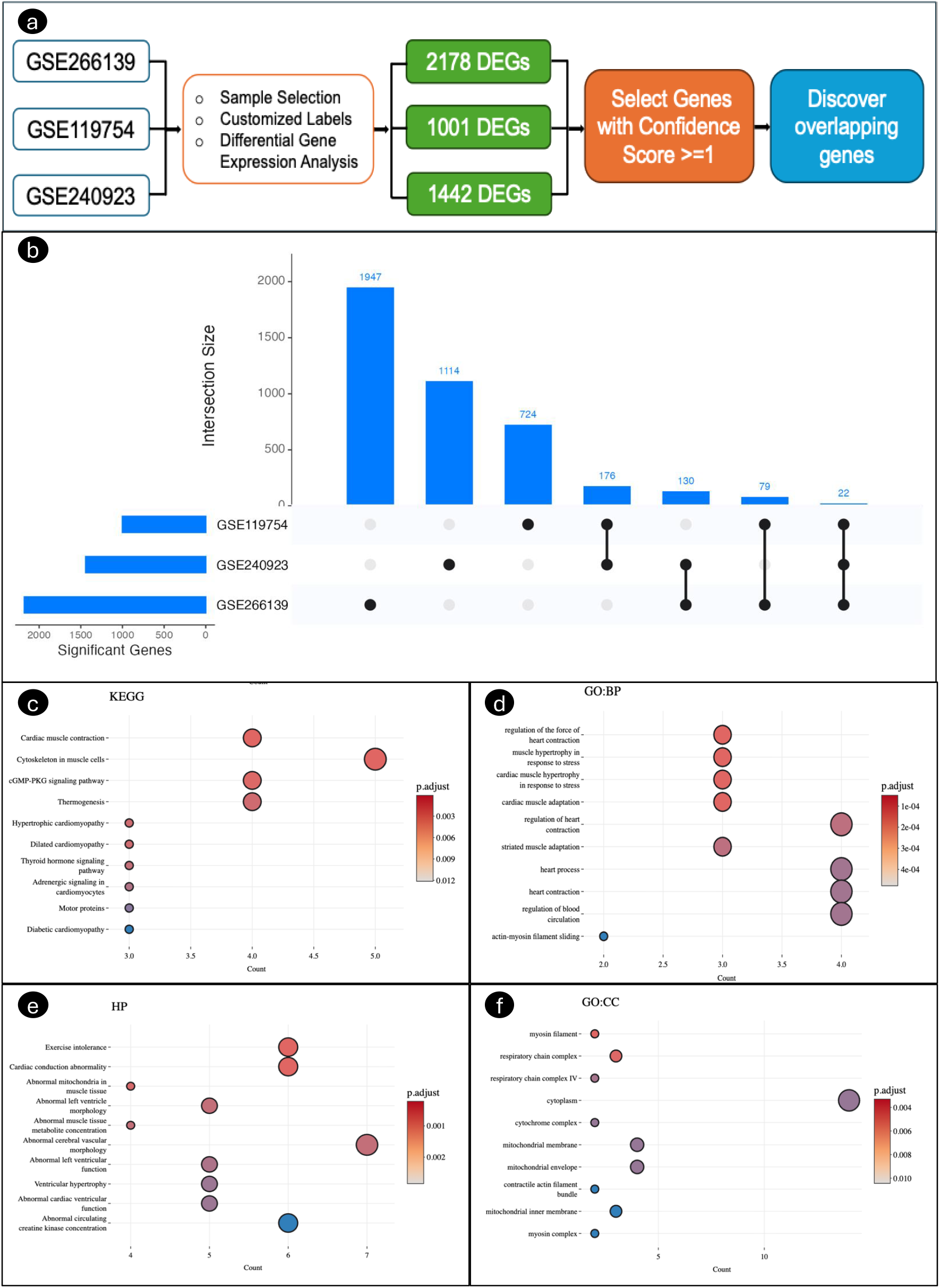
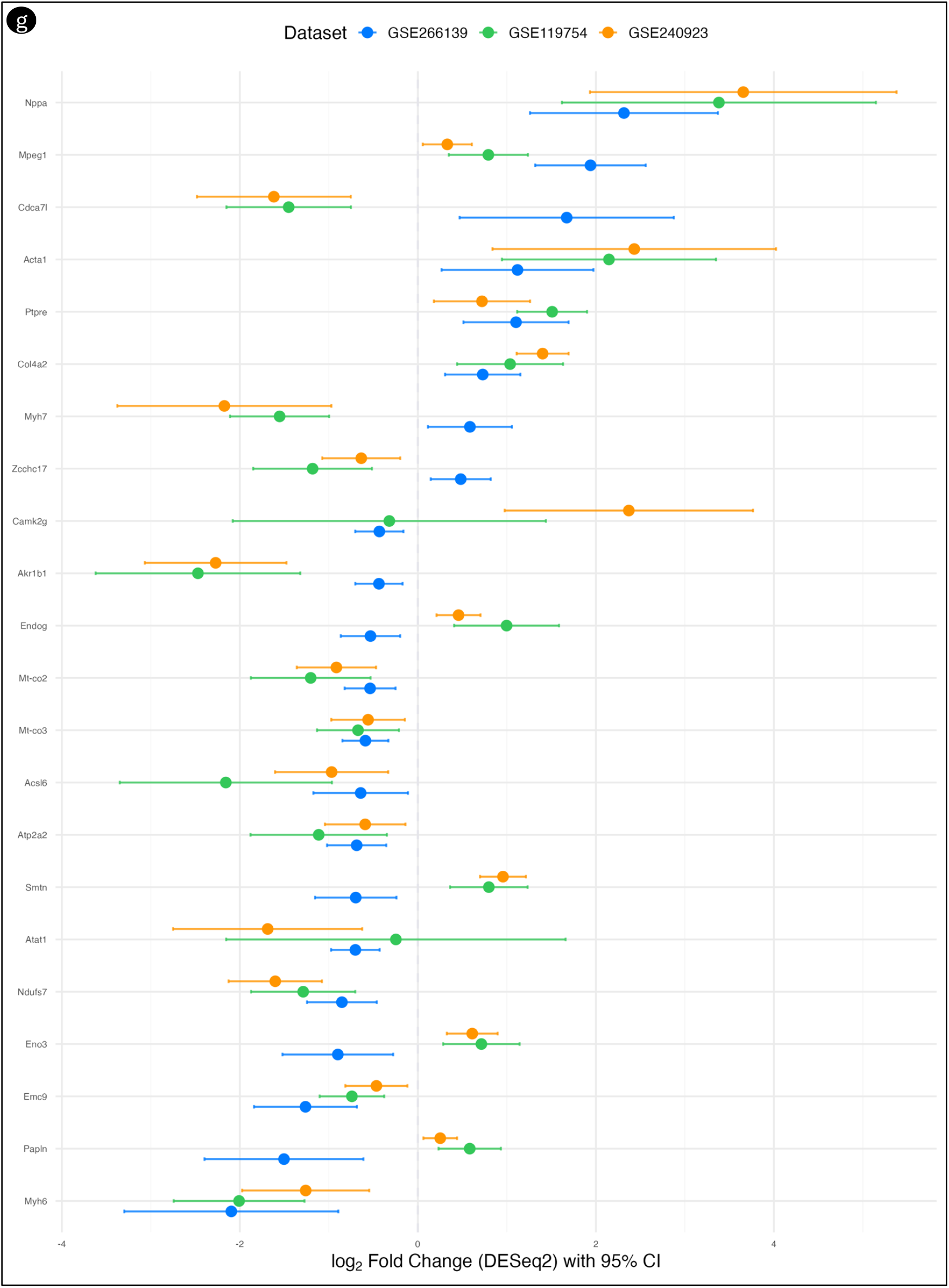
Exploratory meta-analysis across three MCT rat RV datasets. (a) Differential gene expression analysis was performed independently for each GEO dataset, and genes with a Confidence Score ≥ 1 were selected for comparison. (b) UpSet plot showing 22 genes identified as significantly differentially expressed across all three datasets. (c–f) Pathway enrichment analysis of the 22 shared genes. **g)** Forest plot displaying DESeq2 log2 fold-change estimates with 95% confidence intervals for the 22 shared genes across all three datasets, with upregulated genes to the right of zero and downregulated genes to the left. Narrower confidence intervals reflect greater certainty in the estimated change, aiding in gene prioritization

Using PHELEX, we performed an exploratory meta-analysis across three publicly available RNA sequencing datasets profiling right ventricular (RV) tissue in monocrotaline (MCT) pulmonary hypertension rat models: GSE240923 (*19*), GSE119754 (*9*), and GSE266139 (*20*) (**Supplementary data**). Sample-level metadata for each dataset was retrieved and curated within PHELEX, and the experimental characteristics of each dataset are summarized in **Supplementary data**. For each dataset, sample condition labels were assigned through the metadata curation module, gene-by-sample count matrices were assembled, and differential gene expression analysis was performed using the Confidence module with all four analytical methods (DESeq2, edgeR, limma, and NOISeq) (**Figure 5a**).

Across the three datasets, 22 genes were found to be significantly differentially expressed in all three studies with a Confidence Score of at least 1 (**Figure 5b; Supplementary data**). Forest plot showed that several genes were consistently upregulated across all three studies, including *Nppa, Acta1, Ptpre*, and *Col4a2*, while *Mt-co2, Mt-co3, Acsl6, Atp2a2, Ndufs7*, and *Myh6* were consistently downregulated (**Figure 5g**). A subset of genes displayed opposing directionality across the datasets, including *Cdca7l, Myh7, Zcchc17, Emc9, Camk2g, Akr1b1, Endog, Smtn, Eno3*, and *Papln*, potentially due to differences in disease severity, sample collection timepoints, or experimental conditions across the three studies.

Pathway enrichment analysis of the 22 shared genes revealed significant associations with cardiac muscle contraction, cGMP-PKG signaling, hypertrophic cardiomyopathy, and adrenergic signaling in cardiomyocytes (KEGG; **Figure 5c**), as well as cardiac muscle hypertrophy in response to stress, regulation of heart contraction, and regulation of blood circulation (GO:BP; **Figure 5d**). Enriched Human Phenotype Ontology terms included exercise intolerance, ventricular hypertrophy, and abnormal mitochondria in muscle tissue (**Figure 5e**), and GO Cellular Component analysis identified myosin filament and respiratory chain complex terms (**Figure 5f**).

Collectively, these results demonstrate how PHELEX enables rapid identification of biologically coherent and disease-relevant transcriptional signatures across independent PH datasets.

## Discussion

Depending on the platform, modality, and depth of analysis, transcriptomic experiments over the past 15 years have cost between $500 and $1,000 per sample, funds largely derived from public sources. Based on the number of samples we have identified (6,263 unique samples), we estimate that approximately $6.2 million has been spent on generating these data globally. We hypothesize that much of these data have been interrogated a limited number of times, mostly by the originating study, and have not been fully capitalized upon to promote discovery in PH research.

Additionally, datasets available on the Gene Expression Omnibus are difficult to reanalyze (*18, 21*). While the GEO repository is massive, inconsistent metadata and sample annotations make it difficult for researchers to understand and reconstruct the underlying study design. Because of the heterogeneous nature of the studies, and because experimental protocols differ across laboratories, it is difficult to compare datasets in a meaningful way, and deriving biological insights from these data remains challenging (*18*). To address these challenges, we developed PHELEX to democratize access to publicly available PH transcriptomic data for the broader research community.

Here, we present PHELEX for use by the PH community to explore the available data, make useful interpretations, and share results. Through PHELEX, we have acquired raw FASTQ files from GEO, aligned them to the latest species-specific reference genomes, and created uniformly processed count files. Importantly, PHELEX is not simply a data catalogue; by integrating it with Confidence, our differential gene expression analysis module, we offer PH researchers the opportunity to select studies of interest, parse the count files we have generated together with their associated metadata, identify differential gene expression signatures, and place those lists of regulated genes into biological context using pathway enrichment analysis. We have also built in the capacity for users to select multiple studies of interest and compare them, effectively enabling de novo exploratory meta-analysis of PH data, displaying commonly regulated genes through UpSet plots, and visualizing specific genes in forest plots to facilitate high-confidence gene prioritization based on consistent regulation patterns across multiple independent studies.

## Limitations

The application we present here reflects the data that have been uploaded to NCBI’s GEO related specifically to pulmonary hypertension. While we have collected all available data from GEO, the current focus of the tool is solely on RNA sequencing data analysis. Future iterations of PHELEX will provide capacity for the analysis, and hopefully the comparison of multiple data modalities including proteomic, methylomic, and microarray-derived transcriptomic data. PHELEX does not yet have the capacity to analyze or integrate these types of data. At this time, we have provided no functionality for the analysis of single-cell or single-nucleus RNA sequencing data. With the increased availability of this infrastructure, and the lower cost and higher throughput trajectory of the single-cell sequencing ecosystem, we intend to build this functionality into PHELEX in future releases. Furthermore, we currently do not have a mechanism to perform batch correction across datasets, meaning that each dataset must be independently analyzed. Consequently, samples from multiple datasets cannot be truly integrated into a single dataset and analyzed together. Lastly, we currently do not have any machine learning or artificial intelligence (AI) integration, which could change the way that users interact with these data, making hypothesis generation more insight-driven and organic.

## Conclusion

The concept of this project is to empower the entire PH research community with the tools they need to tease biological truth from disparate multi-omic datasets which the PH community has already generated. We hope that this platform will aid target prioritization, target validation, and guide hypothesis generation. We predict PHELEX will have four main outcomes: **1**. That PH scientists will have a single repository within which they can mine to further understand gene expression. **2**. That the gap between different pre-clinical models will shorten, allowing for meaningful comparisons that are linked to haemodynamic or molecular outcomes, **3**. that emerging single cell data is modelled, **4**. that standardized protocols for analysis may emerge, and, **5**. That this platform will be scalable and translate easily to ‘omics data from other disciplines, such as stroke, hypertension, and other types of heart and circulatory diseases. PHELEX is offered without charge to the PH community, and we believe this software will empower PH researchers providing easy access to complex published data that has already been funded; PHELEX will optimize the use of public funds to help find cures for PH.

## Software and source code availability and licensing

The software described in this manuscript, and the source code is protected by copyright. © 2026 Charles Colin Thomas Hindmarch. All rights reserved.

